# A Null Allele of the Pol IV Second Subunit is Viable in *Oryza sativa*

**DOI:** 10.1101/2021.10.21.465363

**Authors:** Tania Chakraborty, Joshua T. Trujillo, Timmy Kendall, Rebecca A. Mosher

**Author notes:** <u>Corresponding author:</u> Rebecca A. Mosher.

## Abstract

All eukaryotes possess three DNA-dependent RNA polymerases, Pols I-III, while land plants possess two additional polymerases, Pol IV and Pol V. Derived through duplication of Pol II subunits, Pol IV produces 24-nt siRNAs that interact with Pol V transcripts to target *de novo* DNA methylation and silence transcription of transposons. Members of the grass family encode additional duplicated subunits of Pol IV and V, raising questions regarding the function of each paralog. In this study, we identify a null allele of the putative Pol IV second subunit, *NRPD2*, and demonstrate that NRPD2 is the sole subunit functioning with NRPD1 in small RNA production and CHH methylation in leaves. Homozygous *nrpd2* mutants have neither gametophytic defects, nor embryo lethality, although adult plants are dwarf and sterile.

## INTRODUCTION

Plant genomes are continuously under threat from transposons, mobile genetic elements whose transposition causes mutation (Dubin, Mittelsten Scheid, and Becker 2018). Host cells use DNA and histone methylation to suppress transposon activity and keep the damage to a minimum. In heterochromatin, transposons are silenced through dense CG methylation and a self-reinforcing loop of non-CG methylation and di-methylation of Histone H3 Lysine 9 (Zemach et al. 2013; Stroud et al. 2014; Du et al. 2012). In gene-proximal regions, transposons are silenced by RNA-directed DNA Methylation (RdDM) (Q. Li et al. 2015; Zhong et al. 2012). Although RdDM can *de novo* methylate cytosines in CG, CHG and CHH contexts (where H is any base other than G), maintenance of CG and CHG methylation by other pathways means that CHH methylation is a primary indication of RdDM (Stroud et al. 2014).

During RdDM, RNA Polymerase (Pol) IV produces short single-stranded RNA transcripts which are simultaneously transcribed into double-stranded RNA by RNA DEPENDENT RNA POLYMERASE 2 (RDR2) (Blevins et al. 2015; Zhai et al. 2015; Singh et al. 2019). These double-stranded RNAs are chopped into 24-nt short interfering (si)RNAs by DICER-LIKE 3 (DCL3), and one of the duplexed siRNAs is loaded into ARGONAUTE 4 (AGO4) (Xie et al. 2004; Kasschau et al. 2007; Singh et al. 2019; Mi et al. 2008; Havecker et al. 2010). AGO4 associates with the carboxy-terminal domain of RNA Pol V, and base-pairing of the nascent Pol V transcript and siRNA recruits DOMAINS REARRANGED METHYLTRANSFERASE 2 (DRM2), which causes *de novo* cytosine methylation (C. F. Li et al. 2006; El-Shami et al. 2007; Wierzbicki et al. 2009; Cao et al. 2003).

Pol IV and V are plant-specific polymerases, which evolved through duplication and functional divergence of multiple Pol II subunits (Luo and Hall 2007; Y. Wang and Ma 2015; Tucker et al. 2010; Huang et al. 2015). Many subunits are shared by Pol II, Pol IV, and Pol V, while unique subunits are presumed to confer the specialized activities of these polymerases (Ream et al. 2009; Haag et al. 2014). In Arabidopsis, the largest subunits (NRPD1 and NRPE1 for Pol IV and V, respectively), are unique, while the second subunit (NRPD2) is shared by Pol IV and Pol V (Ream et al. 2009; Haag et al. 2014). Together, these two largest subunits form the catalytic center of the polymerase. In the grass family, *NRPE1* and *NRPD2* duplicated, giving rise to *NRPF1* and *NRPF2*, respectively (Trujillo et al. 2018). Phylogenetic analysis suggests that these paralogs have new functions, however their functions are not clear.

Mutation of *NRPD1* causes a near-complete loss of 24-nt siRNAs, while disruption of *NRPE1* causes only a small decrease in 24-nt siRNA accumulation (Mosher et al. 2008; Gouil and Baulcombe 2016; Grover et al. 2018; Erhard et al. 2009; Z. Wang et al. 2020; Xu et al. 2020; Debladis et al. 2020; Zheng et al. 2021). However, because both Pol IV and Pol V are required for RdDM, CHH methylation is reduced when either largest subunit is eliminated (Gouil and Baulcombe 2016; Stroud et al. 2013; Zheng et al. 2021). Although Arabidopsis *nrpd1* or *nrpe1* mutants develop normally, loss of *NRPD1* or *NRPE1* in many other species causes developmental defects (Hollick 2010; Gouil and Baulcombe 2016; Grover et al. 2018; Xu et al. 2020; C. Zhang et al. 2020; Z. Wang et al. 2020; Zheng et al. 2021).

Relative to the unique largest subunits, the shared second subunit of Pol IV and V is less studied. Arabidopsis *nrpd2* mutants lack 24-nt siRNAs and CHH methylation and do not accumulate Pol V transcripts (Herr et al. 2005; Kanno et al. 2005; Onodera et al. 2005; Pontier et al. 2005; X. Zhang et al. 2007; Wierzbicki, Haag, and Pikaard 2008). Like mutations of the largest subunits, *nrpd2* plants do not exhibit developmental phenotypes in Arabidopsis. In maize, there are three second subunit paralogs (two copies of NRPD2 and one copy of NRPF2) and phylogenetic analysis suggests these paralogs are functionally distinct (Haag et al. 2014; Trujillo et al. 2018). Mutants of *NRPD2a* show disrupted 24-nt siRNA production and defective paramutation, suggesting that there is little redundancy between paralogs (Sidorenko et al. 2009; Stonaker et al. 2009), while immunoprecipitation of polymerase complexes also hints that the duplicated NRPD2 subunits in grasses are not functionally redundant (Haag et al. 2014; Trujillo et al. 2018).

RdDM is critical for development in rice. Loss of *NRPD1* causes sterility, reduced plant height, and increased tillering (Xu et al. 2020; C. Zhang et al. 2020; Zheng et al. 2021). Mutations in other components of the RdDM pathway in rice have similar developmental phenotypes. *dcl3* RNAi lines showed reduced plant height and increased flag leaf angle, along with reduced panicle size (Wei et al. 2014). *ago4ab* and *rdr2* RNAi lines also showed reduced height and panicle size, and the *ago4ab* line phenocopied *dcl3* in flag leaf angle (Wei et al. 2014). *drm2* mutants showed a wide range of developmental phenotypes including complete sterility (Moritoh et al. 2012).

In this study, we characterize a rice T-DNA insertion mutant in *NRPD2*, the gene presumed to encode the second subunit for both Pol IV and Pol V. We show that the *nrpd2* mutant loses nearly all 24-nt siRNAs, especially from TE-rich regions, and also loses CHH methylation in specific sites. These molecular and developmental phenotypes largely copy *nrpd1* mutants, indicating that *NRPD2* is the sole second subunit for Pol IV in leaves.

## RESULTS

### Rice *NRPD2* is required for 24-nt siRNA production

To investigate the function of NRPD2 in rice, we searched the PFG T-DNA insertion library and obtained a line with an insertion in the Rpb6 domain (**Figure 1A**) (Jeon et al. 2000; Jeong et al. 2006). RT-PCR confirmed the loss of *NRPD2* gene expression in plants homozygous for the mutation (**Figure 1B**). Sequencing wild-type small RNA populations confirmed that 24-nt small RNAs are the plurality of total small RNAs in *Oryza sativa* cv. Japonica var. Hwayoung, totaling 46% of small RNAs in leaves. In comparison to wild type, *nrpd2* mutants show ~69% reduction of 24-nt siRNAs (**Figure 1C**). This reduction is consistent with *nrpd2* mutants in Arabidopsis (X. Zhang et al. 2007), and *nrpd1* mutants in rice (Xu et al. 2020; Debladis et al. 2020; Zheng et al. 2021). 21- and 22-nt sRNAs are elevated in *nrpd2* populations, most likely due to oversampling of these sizes following depletion of the dominant 24-nt size class. Together, these results show that *NRPD2* is required for majority of the 24-nt small RNA production in rice leaves.

**Figure 1.**
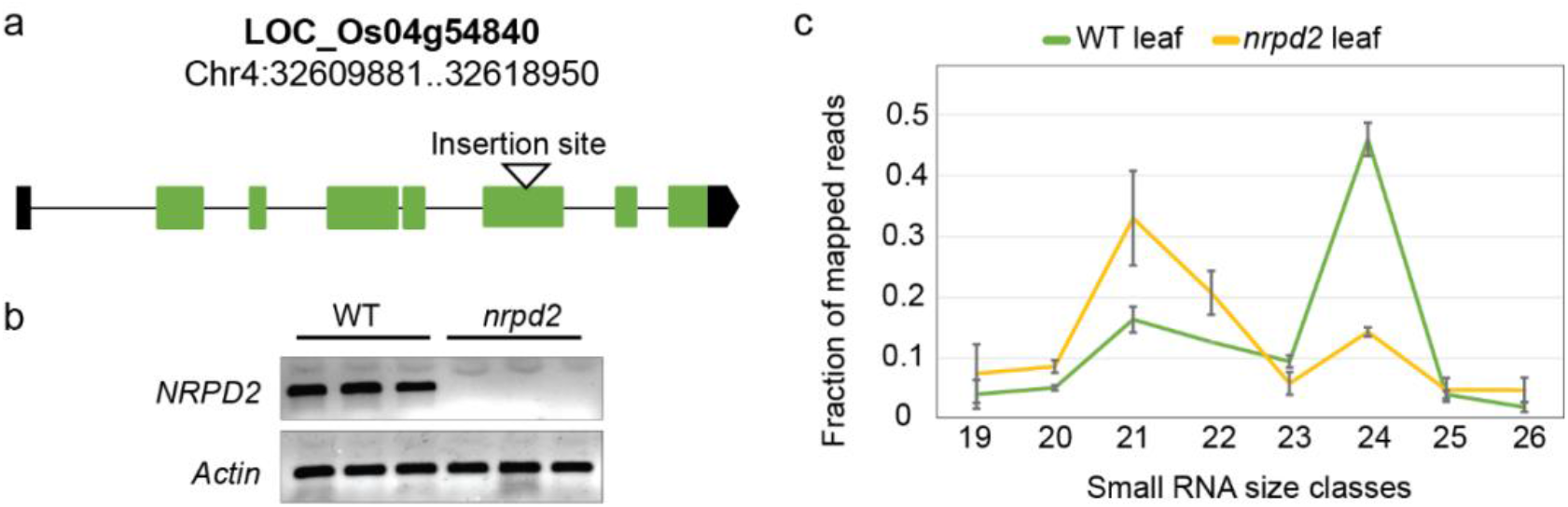
A T-DNA insertion in *NRPD2* gene results in loss of 24-nt small RNAs. **(a)** Schematic of T-DNA insertion in the *Oryza sativa NRPD2* gene. Green boxes indicate coding exons, black boxes depict untranslated regions. The T-DNA insertion site is marked with a white triangle. **(b)** RT-PCR of *NRPD2* and *ACTIN* in wild-type (WT) and *nrpd2* mutant leaves. Three independent biological replicates are shown for each genotype. **(c)** Line graph showing size profiles of filtered and mapped 19-26 nt small RNAs from wild type (WT) and *nrpd2* mutants. The average of three independent biological replicate libraries is shown; error bars show standard deviation.

### NRPD2 function overlaps sites of NRPD1 action

To directly compare the function of NRPD2 with NRPD1, we obtained sRNA sequencing data from rice *nrpd1a nrpd1b* double mutants (*nrpd1*, hereafter) (Zheng et al. 2021) and used the 300-bp non-overlapping windows to count small RNAs. There was a good correlation between wild-type sRNA accumulation in leaf and seedling tissues (**Figure 2A**). Because there is 24-nt siRNA depletion in both *nrpd2* and *nrpd1* mutants, we wanted to understand if this loss was distributed across many genomic loci, or if it came from a small subset of abundantly-expressed loci (Burgess et al. 2021). To examine this, we assessed sRNA accumulation in 300-bp windows. Among windows with at least 5 mapped reads in wild type in any size class, those with less than 10% of wild-type sRNA accumulation were called “depleted windows”, while windows showing an increase in sRNA accumulation compared to wild type were called “independent windows”. We observed that in the *nrpd2* mutant there are 188,215 depleted windows (~78%), indicating that the loss of small RNAs observed in the mutant occurs at most small RNA-producing loci. We observed a similar percentage of siRNA-depleted windows (~77%) in *nrpd1* (**Figure 2B**).

**Figure 2.**
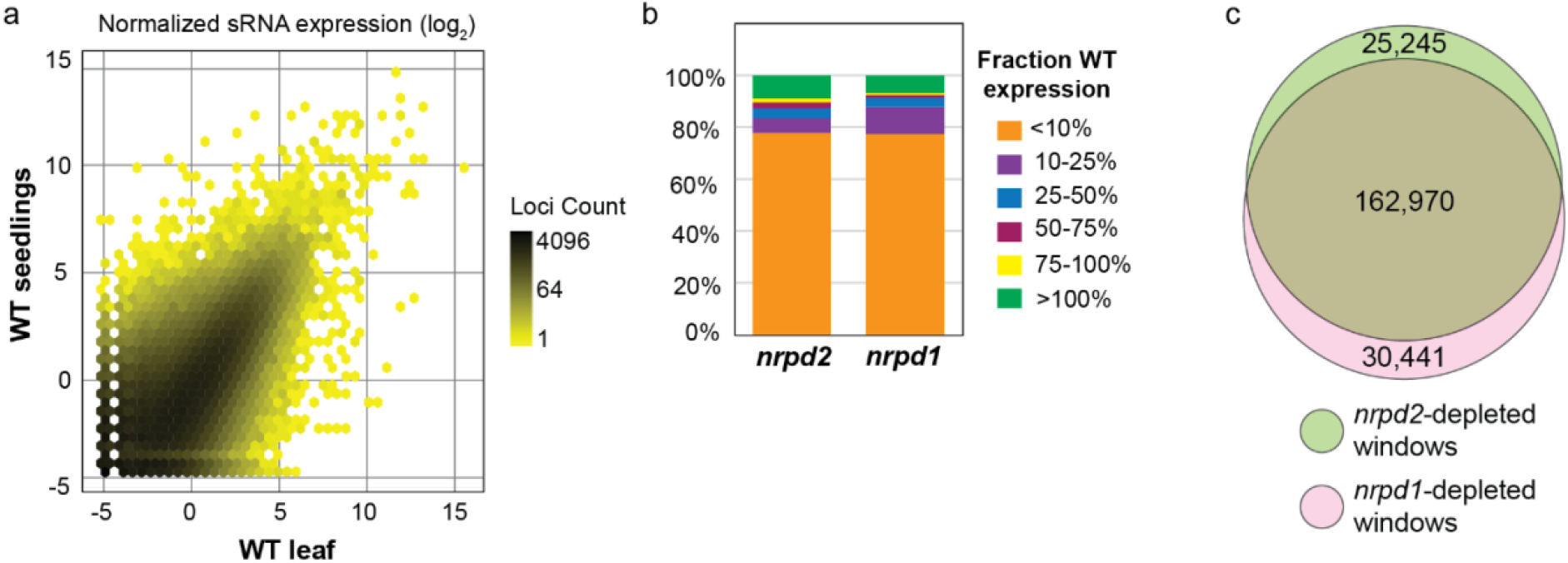
*nrpd2* mutants lose small RNAs from same locations as *nrpd1*. (a) Correlation plot with small RNA levels in wild-type leaves and wild-type seedling. (b) Stacked bar graph showing normalized expression levels in *nrpd2* and *nrpd1* relative to wild-type expression levels. (c) Venn diagram showing proportion of overlap between *nrpd2*-depleted and *nrpd1*-depleted windows. Windows were designated as “depleted” if small RNA accumulation was reduced to ≤10% of wild type. *nrpd1* small RNA data is from (Zheng et al., 2021).

To better understand the genomic regions where NRPD2 acts, we examined overlaps between *nrpd2*-depleted or -enriched windows and genomic features like genes, transposons, miRNAs, centromeric and telomeric repeats. We observed that the *nrpd2*-depleted windows were significantly enriched for DNA transposons, MITE elements, and SINE retrotransposons, but were depleted in genes and LTR retrotransposons (**Table 1)**. This pattern is similar to regions enriched or depleted at sites of RdDM in *Brassica rapa* (Grover et al. 2018), indicating that *nrpd2*-depleted windows resemble canonical RdDM loci. The *nrpd2*-independent windows overlapped significantly with genes, miRNAs, and short transposons (MITEs and SINEs). To understand if the overlap of *nrpd2*-independent windows with MITEs was because of the physical proximity of the MITEs and SINEs to genes, which accumulate *nrpd2*-independent small RNAs, we calculated the distance between genes and the MITEs and SINEs that overlap the *nrpd2*-depleted and *nrpd2*-independent windows. We observed that while most *nrpd2*-depleted windows that overlap MITEs and SINEs are far away from genes, majority of *nrpd2*-independent windows that overlap MITEs and SINEs also overlap genes (**Supplemental Figure 1**). These observations demonstrate that *nrpd2*-depleted windows resemble canonical RdDM loci.

**Table 1:** Enrichment of genomic features across *nrpd2*-depleted and *nrpd2*-independent windows

| Feature | Fold change |  |
| --- | --- | --- |
|  | *nrpd2*-depleted windows | *nrpd2*-independent windows |
| DNA Transposons | 1.9*** | n.s. |
| MITE Transposons | 2.3*** | 2.1*** |
| LTR Transposons | 0.52*** | 0.73*** |
| LINE Transposons | 0.83* | 0.24** |
| SINE Transposons | 9.6*** | 7.4*** |
| Telomere repeats | 16.5*** | n.s. |
| Genes | 0.44*** | 5.2*** |
| lmiRNAs | n.s. | 61.1** |
| miRNAs | 0.10*** | 25.3*** |
| Centromeric repeats | n.s. | n.s. |

\*, $p < 0.05$ , \*\*, $p < 10^{-4}$ , \*\*\*, $p < 10^{-10}$

To investigate whether *nrpd2* affected the same genomic regions as a loss of *nrpd1*, we overlapped the small RNA depleted windows from both mutants. We observed a significant overlap between the two sets of genomic locations (**Figure 2C**) and a similar enrichment of SINE and MITE elements in *nrpd1*-independent windows, indicating that NRPD1 and NRPD2 are required in the same locations.

Together, these data indicate that *nrpd2* phenocopies *nrpd1* in rice leaf, suggesting that NRPD2 is the sole second subunit assembled in Pol IV in leaves.

### Rice NRPD2 participates in RdDM

To confirm that the siRNAs produced by NRPD2 act in RdDM, we assayed methylation with quantitative Chop-PCR at loci producing *nrpd2*-dependent 24-nt siRNAs (H. Zhang et al. 2014). Loci were randomly selected based on the presence of an *NIa III* site, at least 90% reduction in 24-nt siRNA accumulation in the *nrpd2* mutant, and low variability in qPCR. All 15 of the windows showed higher methylation in wild type than *nrpd2*, although not all changes are significant and sometimes methylation in wild type was low, perhaps as a consequence of the assay testing methylation at a single cytosine (**Supplemental Figure 2**). Of the 8 windows with at least 5% methylation in WT, 7 were substantially reduced in *nrpd2* (**Figure 3**). These results indicate that CHH methylation at *nrpd2*-depleted windows requires NRPD2.

**Figure 3.**
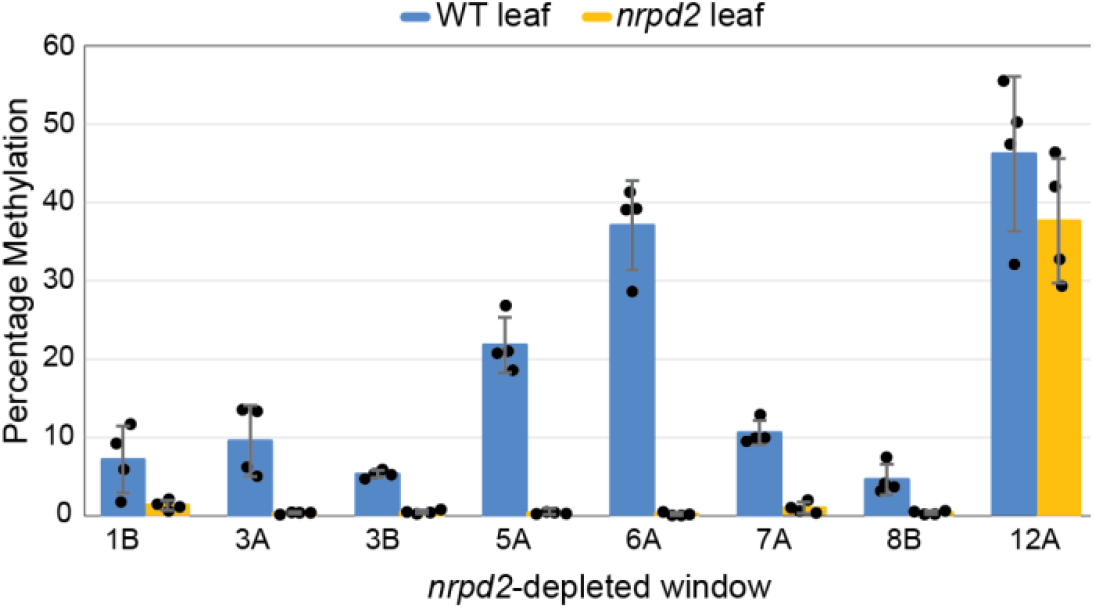
*NRPD2* is required for RdDM. Quantitative methylation-sensitive PCR (chop-qPCR) for CHH methylation at *nrpd2-*depleted windows demonstrates that *NRPD2* is required for RdDM. The mean and standard deviation of four independent biological replicates are shown in bars. Individual replicates are shown as black dots.

### *nrpd2* mutants are viable but infertile

Rice lacking Pol IV or Pol V function show reduced height and drastic loss in fertility (Zheng et al. 2021). To investigate the morphological phenotypes of the *nrpd2* mutant, we grew selfed seeds from a heterozygous plant and observed 11 wild type, 23 heterozygotes, and 9 homozygous mutants, indicating there were neither gametophytic nor embryonic defects associated with loss of *NRPD2* function. Heterozygous *nrpd2* mutants were indistinguishable from their wild-type siblings, however homozygous *nrpd2* mutants were substantially shorter (**Figure 4A**). The homozygous *nrpd2* mutants also produced fewer tillers than their wild-type and heterozygous siblings (**Figure 4B**), in contrast to *nrpd1*, which is reported to increase tiller number (Xu et al. 2020). This difference might be due to different genetic backgrounds, as the *nrpd1* mutation is in the Japonica cultivar Nipponbare, while the *nrpd2* mutation is in the Japonica cultivar Hwayoung. Although homozygous *nrpd2* mutants produced panicles, these were small and frequently did not emerge from the sheath (**Figure 4C, D**). Florets on these panicles were misshapen and never produced viable seed, indicating that *NRPD2* function is essential for reproductive development.

**Figure 4.**
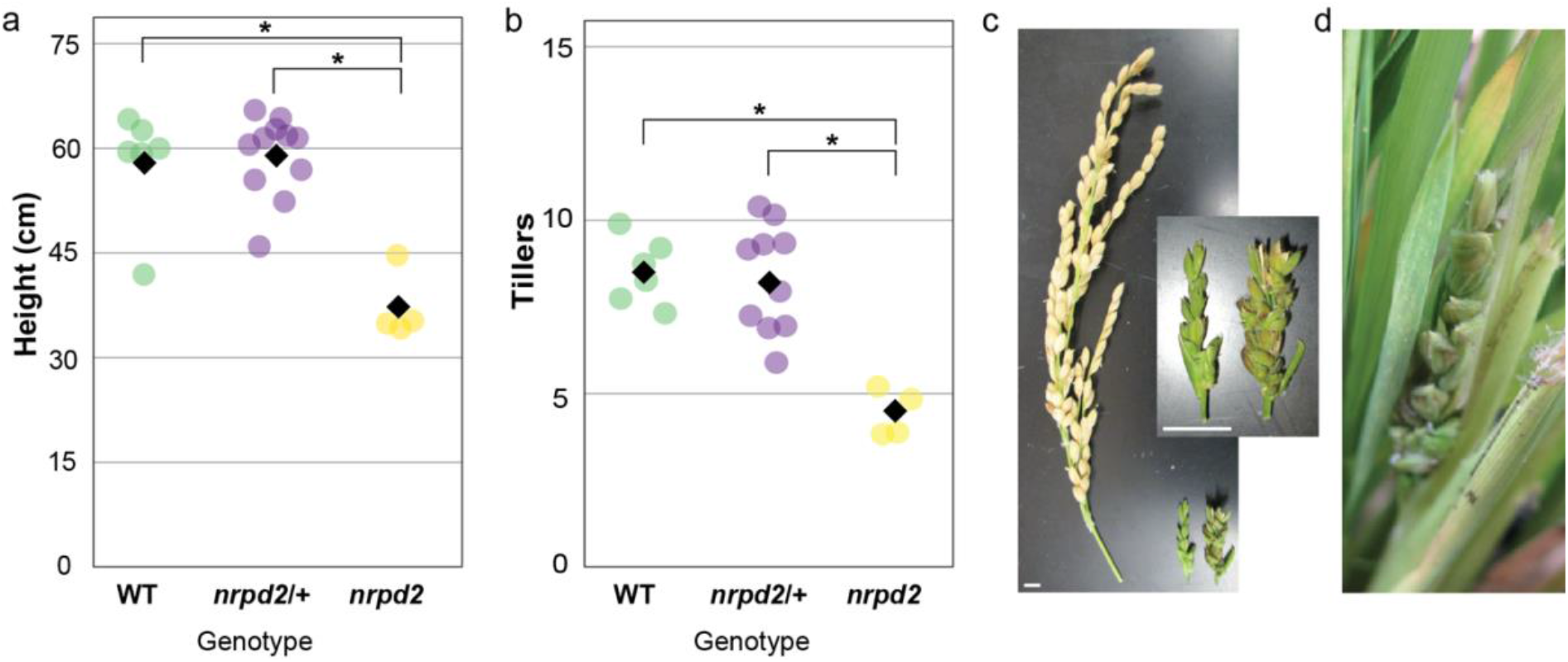
Developmental defects in the *nrpd2* mutant. (a) Jitter plot showing height of wild type, heterozygous, and homozygous *nrpd2* mutants. (b) Jitter plot showing tiller number in wild type, heterozygous, and homozygous *nrpd2* mutants. Mean of measurements shown as black diamonds. One way ANOVA was performed to test significance, * depicts p < 0.005. (c) Panicles produced in wild type (left) or the *nrpd2* mutant (right and inset). Scale bars, 1 cm. (d) Example of an *nrpd2* panicle stuck in the sheath.

## DISCUSSION

RNA-directed DNA methylation silences euchromatic transposons, potentially influencing the expression of nearby genes (Dubin, Mittelsten Scheid, and Becker 2018). Although the rice genome is smaller than many flowering plants, approximately half of its genome is transposons, including a large number of MITEs (Minature Inverted repeat Transposable Elements) near genes (Jiang and Panaud 2013; Song and Cao 2017). Thus, RdDM is expected to be important both for transposon control and gene expression in rice. As rice is one of the important food crops across the world, it is therefore essential to understand the mechanism of RdDM in rice. We characterized a T-DNA insertional mutation in *Oryza sativa NRPD2*, the putative second subunit of RNA Pol IV and Pol V, and provided evidence that this subunit is required for Pol IV function.

Poaceae-specific duplication of putative Pol V largest (NRPE1) and second-largest (NRPD2) subunits raises interesting questions about the assembly of multiple non-canonical polymerases in rice (Trujillo et al. 2018). Zheng and colleagues recently demonstrated that NRPF1 is not redundant with NRPE1, and not required for RdDM in seedlings (Zheng et al. 2021), suggesting that each paralogous largest subunit has unique function. However, sharing of the second subunit between Arabidopsis Pol IV and Pol V suggests that Pol function is primarily determined by the largest subunit and that second subunits might be redundant. We have shown that *nrpd2* mimics *nrpd1* with respect to small RNA accumulation, DNA methylation, and developmental phenotype. We therefore conclude that NRPD2 is the sole second subunit assembled in Pol IV and is not redundant with NRPF2. While our molecular analysis is limited to Pol IV activity in leaves, shared developmental phenotypes indicate that NRPD1 and NRPD2 function together exclusively throughout rice development.

NRPD2 targets are enriched for transposons, including MITEs, which are frequently found in proximity to genes in rice (Jiang and Panaud 2013). Loss of methylation at MITEs in the *nrpd1* mutant causes increased expression of a miRNA gene and downregulation of its target protein-coding gene, ultimately causing increased tillering (Xu et al. 2020). We did not observe increased tillering in *nrpd2* mutants; indeed, homozygous mutants have fewer tillers than wild-type or heterozygous siblings, and more closely resemble knockdown mutants of *dcl3* or *drm2* (Moritoh et al. 2012; Wei et al. 2014). Embryonic lethality was also reported for *nrpd1* mutants, however, we observe homozygous mutant seeds arising at the expected ratios, indicating there is no lethality for homozygous mutant embryos. *NRPE1* mutations in different rice strains result in different phenotypes (Zheng et al. 2021), suggesting that genomic differences between rice strains, especially with respect to transposon insertion site, might explain the differences between *nrpd1* and *nrpd2* phenotypes we observed.

A phenotype shared by all of the various rice RdDM mutations is reduced fertility (Moritoh et al. 2012; Wei et al. 2014; Xu et al. 2020; Zheng et al. 2021). RdDM is also required for full fertility in a range of other angiosperms (Chow, Chakraborty, and Mosher 2020), suggesting that defects in reproduction are deeply conserved and less likely to result from transposon-mediated regulation of individual genes. Given the importance of rice reproduction for the global food supply, understanding how genes like *NRPD2* contribute to successful reproduction is a critical area of research.

## METHODS

### Plant Material and Growth Conditions

The PFG T-DNA allele of *NRPD2* (PFG_1A-24731.R) is in the *Oryza sativa* (Japonica) Hwayoung background. All plants were grown in a greenhouse at 85°F and were genotyped with primers flanking the T-DNA insertion site and T-DNA border primers (**Supplemental Table 1**). Wild type amplifies a product only with the flanking primers, while homozygous *nrpd2* mutants amplify products with a combination of flanking and border primers. Heterozygous mutants amplify products with both combinations of primers. Homozygous plants and wild-type sibling controls were derived from self-fertilized heterozygous parents.

### Small RNA sequencing

Leaf tissue was collected from three wild type and three homozygous *nrpd2* mutant siblings and total nucleic acid was prepared (White and Kaper 1989). The small RNA (<200 nt) fraction was purified with the mirVana miRNA isolation kit (ThermoFisher) prior to small RNA sequencing library preparation using the NEBNext Small RNA kit (NEB; E7330). Libraries were pooled and sequenced by The University of Arizona Genetics Core on an Illumina NextSeq500. These datasets are available in the National Center for Biotechnology Information Sequence Read Archive under Bio Project PRJNA758109.

### Small RNA analysis

FastQC (Andrews 2010) was used to quality-check the raw small RNA reads that were obtained from the sequencing facility. Adapters were removed with Trim Galore, with options - length 10 and -quality 20 (Krueger v0.6.5 2019). After that, reads mapping to the *Oryza sativa, Brassica rapa*, and *Arabidopsis thaliana* chloroplast and mitochondrial genomes were removed by aligning to these genomes using Bowtie (Langmead et al. 2009). Reads mapping to structural and noncoding RNAs were also removed by aligning to the rfam database (Kalvari et al. 2018) using all available *Oryza sativa* sequences. After this, reads at least 19-nt and up to 26-nt were selected and aligned to the *Oryza sativa* var. Japonica genome using Bowtie. Only sRNA reads with a perfect genomic match were kept for later analysis. ShortStack (Axtell 2013) was used to annotate all small RNA loci, with options-mismatches 0, -mmap u, -mincov 0.5 rpm, and -pad 75. All of these steps can be done with a small RNA analysis pipeline (Bose 2019). ShortStack was used again to count small RNAs on 300-bp tiled windows across the genome. Upon investigation of the windows, it was observed that four windows produced a large number of small RNAs, almost to a tenth of the whole library size. Upon a BLAST search, we observed that most of the small RNAs mapped to rRNAs in the Rfam database, and therefore we removed all the reads from these four windows. The reads were normalized to reads per million mapped reads (RPM) and all further analysis were performed on normalized reads. The number of reads at each step in each replicate can be found in **Supplemental Table 2**.

Principal component analysis was done on siRNA accumulation in each 300-bp window to check for consistency of the replicates (**Supplemental Figure 3**), and size profiles of mapped small RNAs in the replicates was used as an additional check.

To measure the impact of the *nrpd2* mutation, only windows whose expression across 19-26mers was at least 0.5 RPM in both our leaf wild-type and seedling wild-type (Zheng et al. 2021) samples were considered for further analysis. *rpd2*-depleted and *nrpd2*-independent windows were overlapped with genomic features using “bedtools overlap” function (Quinlan and Hall 2010) and overlaps of at least 1 nt were counted. Expected values were calculated based on the number of 300-nt tiled windows containing the genomic features and Fisher’s exact tests were performed to understand the significance of enrichment or depletion. Only significant overlaps are reported.

For the correlation plot, we plotted normalized small RNAs across windows in both wild-type samples. For the Venn diagram, we obtained the list of “*nrpd2*-depleted” and *“nrpd1-* depleted” windows and calculated the overlap between those with “bedtools overlap” function. Then we used the Venn Diagram Plotter (Kyle Littlefield and Matthew Monroe 2020).

### RT-PCR and Chop-qPCR for methylation analysis

For RT-PCR of *ACTIN* and *NRPD2*, total nucleic acid was prepared as described above and treated with DNase using the DNA-free DNA Removal Kit (ThermoFisher). A One-step RT-qPCR kit (Promega) was used for converting the RNA into cDNA before amplifying target transcripts on a CFX Connect Real-Time PCR Detection System (Biorad). RT-PCR primers are listed in **Supplemental Table 1**.

For Chop-qPCR, we gathered the genomic sequence of windows where *nrpd2* and *nrpd1* showed at least 90% reduction in 24-nt siRNAs compared to wild type and an *Nla III* recognition sequence (CATG) was present. *Nla III* is sensitive to methylation of the first C in its recognition sequence, a CHH site. We digested 1 μg of wild type or *nrpd2* DNA with *Nla III* and also performed a “mock” digestion without enzyme. Then we used the digested and mock treated samples to amplify these twenty-five loci as well as *ACTIN* as a control. Amplification was quantified with a CFX Connect Real-Time PCR Detection System (Biorad) qPCR machine.

Relative percentage methylation at each site was calculated by comparing the Cq values for digested and mock digested treatments for each sample (H. Zhang et al. 2014). Chop-qPCR primers are listed in **Supplemental Table 1**.

## Supplemental Information

Supplemental Figures 1-3, Supplemental Table 1-2

## Authors’ contributions

RAM conceptualized the project; TC, TK, and JT performed experiments; TC and RAM analyzed and visualized the data and wrote the paper.

## Acknowledgements

The authors gratefully acknowledge support from the National Science Foundation (MCB-1929678 to RAM). The authors would also like to extend gratitude to Dr. Jesse Woodson for use of his qPCR machine, and Ms. Kyungsook An for sending the PFG T-DNA lines. Illumina sequencing was performed by The University of Arizona Genetics Core.

