## Supplemental Materials for "A Null Allele of the Pol IV Second Subunit is Viable in *Oryza sativa*"

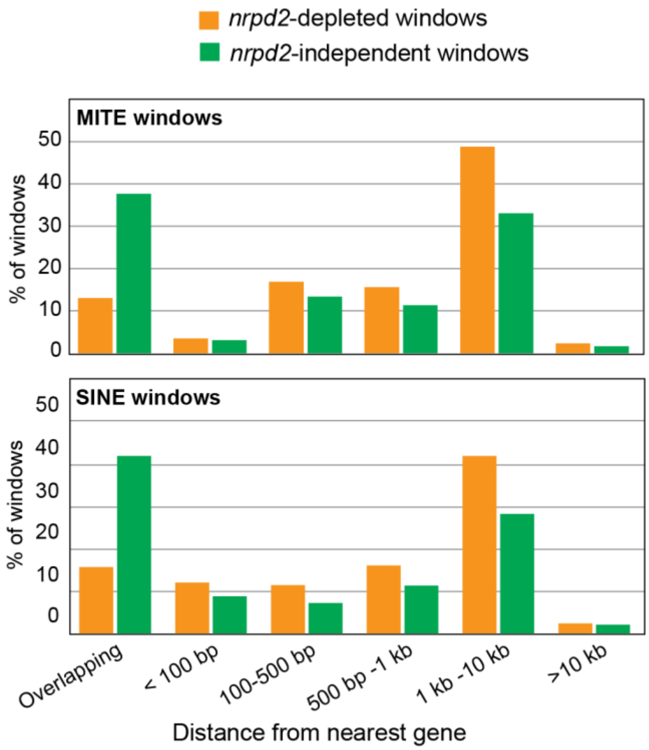


**Supplemental Figure 1**. Percentage of *nrpd2*-depleted and *nrpd2*-independent windows overlapping with MITE transposons that are at each category of distance from the nearest genes

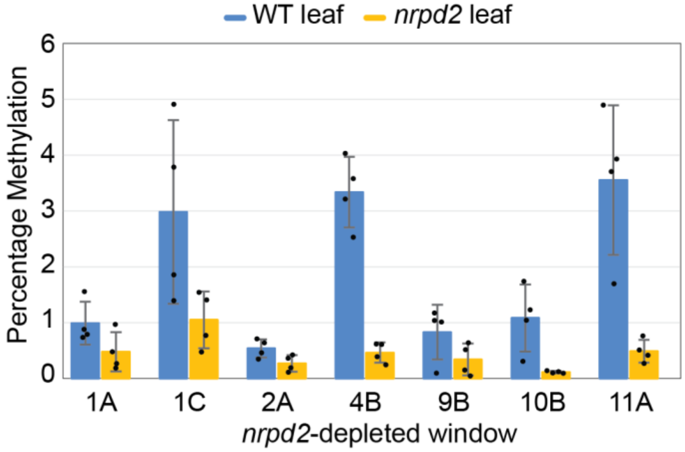


**Supplemental Figure 2.** Chop-qPCR at loci where WT methylation < 5%.

The mean and standard deviation of four independent biological replicates are shown in bars. Individual replicates are shown as black dots.

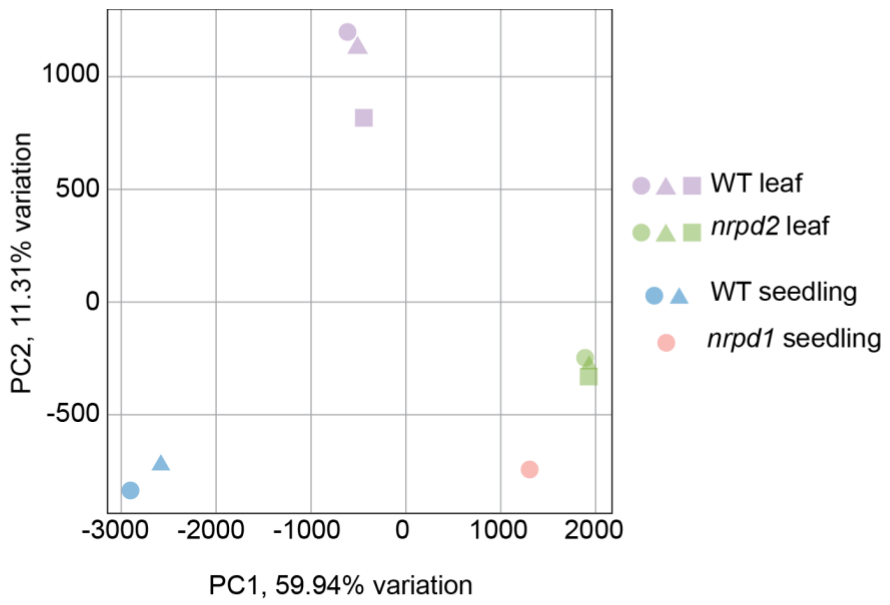


**Supplemental Figure 3.** Principle Component Analysis (PCA) of sRNA libraries, based on small RNA accumulation in 1,244,155 300-nt windows. Seedling samples from Zheng et al. 2021.

**Supplementary Table 1.** Primers used in the study

| Name | Sequence |
|---|---|
| **Genotyping** | |
| PFG_1A-24731.R GT RP | CCAATGTCAGGATCATGAATATGCTG |
| PFG_1A-24731.R GT LP | GTGTTTTCTGATGCAGGTCGAATC |
| pGA2707 RB1 | CCACAGTTTTCGCGATCCAGACTG |
| pGA2707 RB2 | TTGGGGTTTCTACAGGACGTAAC |
| pGA2707 LB1 | GGTGAATGGCATCGTTTGAA |
| pGA2707 LB2 | ACAAGCCGTAAGTGCAAGTG |
| **RT-PCR primers** | |
| actin_rice_cds_f | GCAACTGGGATGATATGGAGAAGA |
| actin_rice_cds_r | TGAAGGTCTCAAACATGATCTGGG |
| nrpd2_cds_f | GTGTTTTCTGATGCAGGTCGAATC |
| nrpd2_cds_r | TATTCTGATCTACCAATGCAGTCAG |
| **Chop-PCR** | |
| 1A_F | AGCTATATTCTCGGAACTTTGGGG |
| 1A_R | GCTTTCAACACTTGGGTCTGAATT |
| 1B_F | AGTGGATGACAATGATACTGTGAT |
| 1B_R | TCCATATGCTTCCTTTTGCTCATG |
| 1C_F | TAAGTCTCTTCCCTCCTCATTCGA |
| 1C_R | ATGCCAACCCTAATAGCTCTCTAC |
| 2A_F | ATTGAATCTGATGGTTGCGTACTT |
| 2A_R | AAGCGGAGTTGTTCTTCTTTCTTG |
| 3A_F | AATACAATAAGTGACCTGCAACGC |
| 3A_R | ATAGGAGAAACACGTCAGCTACAG |
| 3B_F | CAGGTATTGTGGAGGTAGTAAGCA |
| 3B_R | TATTGGAGGTAATAAGATGGGTCCC |
| 4B_F | GAACGAAACCGAAGAGCCATATTC |
| 4B_R | ACAAGTGACCAATTTTAGCTACGC |
| 5A_F | AAATGTACTTACCCTCACGAGCAT |
| 5A_R | AATTTTACTACACTTGCGAGCCAA |
| 6A_F | TCGATCCCTTCTCCTGTACTACTT |
| 6A_R | CCAATTTGTGAAAATGCGGTTTGA |
| 7A_F | AGCATACAAGTGGACTAGGAATCA |
| 7A_R | GGTAACATTGTAGCCTGCTTACAT |
| 8B_F | ACATGTGTTTCCTGTCCAGATTCT |
| 8B_R | ACAGTGCATAGCTAGTGAATCTCA |
| 9B_F | TGATAGCACAAGGGTCATTTCTCT |
| 9B_R | TGAATCGAGCAAAAGGTTCCAATC |
| 10B_F | TTCGTGGGTGTGTAAGTATAACGT |
| 10B_R | ACCCACGAATAAGCCCATAACATA |
| 11A_F | CGATGGTATGATTGTGTTGTAAGCA |
| 11A_R | TTATGCCACCCCGATCTATTTGAC |
| 12B_F | CGATGACTAGTAACACGGGATATC |
| 12B_R | CCCGAGCAATGTGAATCTAGGAAA |

**Supplementary Table 2.** Small RNA sequencing data

| Genotype, Replicate | NCBI identifier | De-multiplexed Reads | Adapter and q>30 Trimming | % Remaining after trimming | Chloroplast, mitochondria, and rfam filtering | % Remaining of trimmed reads after [illegible] | Reads mapped to Nipponbare | Filtering additional structural RNAs | % Trimmed reads analyzed |
|---|---|---|---|---|---|---|---|---|---|
| WT Leaf, 1 | SRX11930857 | 61,052,596 | 21,615,676 | 35.4% | 14,984,953 | 69.3% | 4,757,891 | 4,751,330 | 22.0% |
| WT Leaf, 2 | SRX11930858 | 49,910,517 | 20,223,704 | 40.5% | 13,881,036 | 68.6% | 4,420,133 | 4,411,832 | 21.8% |
| WT Leaf, 3 | SRX11930859 | 69,105,942 | 20,016,003 | 29.0% | 13,933,718 | 69.6% | 6,487,852 | 6,483,749 | 32.4% |
| nrpd2 Leaf, 1 | SRX11930854 | 49,579,598 | 15,487,264 | 31.2% | 9,517,114 | 61.5% | 2,323,033 | 2,316,967 | 15.0% |
| nrpd2 Leaf, 2 | SRX11930855 | 55,632,528 | 19,757,935 | 35.5% | 10,595,617 | 53.6% | 1,862,219 | 1,854,240 | 9.4% |
| nrpd2 Leaf, 3 | SRX11930856 | 61,259,353 | 11,740,487 | 19.2% | 6,568,221 | 55.9% | 2,952,740 | 2,948,756 | 25.1% |
